# Computational Mechanobiology Model Evaluating Healing of Postoperative Cavities Following Breast-Conserving Surgery

**DOI:** 10.1101/2023.04.26.538467

**Authors:** Zachary Harbin, David Sohutskay, Emma Vanderlaan, Muira Fontaine, Carly Mendenhall, Carla Fisher, Sherry Voytik-Harbin, Adrian Buganza Tepole

## Abstract

Breast cancer is the most commonly diagnosed cancer type worldwide. Given high survivorship, increased focus has been placed on long-term treatment outcomes and patient quality of life. While breast-conserving surgery (BCS) is the preferred treatment strategy for early-stage breast cancer, anticipated healing and breast deformation (cosmetic) outcomes weigh heavily on surgeon and patient selection between BCS and more aggressive mastectomy procedures. Unfortunately, surgical outcomes following BCS are difficult to predict, owing to the complexity of the tissue repair process and significant patient-to-patient variability. To overcome this challenge, we developed a predictive computational mechanobiological model that simulates breast healing and deformation following BCS. The coupled biochemical-biomechanical model incorporates multi-scale cell and tissue mechanics, including collagen deposition and remodeling, collagen-dependent cell migration and contractility, and tissue plastic deformation. Available human clinical data evaluating cavity contraction and histopathological data from an experimental porcine lumpectomy study were used for model calibration. The computational model was successfully fit to data by optimizing biochemical and mechanobiological parameters through the Gaussian Process. The calibrated model was then applied to define key mechanobiological parameters and relationships influencing healing and breast deformation outcomes. Variability in patient characteristics including cavity-to-breast volume percentage and breast composition were further evaluated to determine effects on cavity contraction and breast cosmetic outcomes, with simulation outcomes aligning well with previously reported human studies. The proposed model has the potential to assist surgeons and their patients in developing and discussing individualized treatment plans that lead to more satisfying post-surgical outcomes and improved quality of life.

## 1. Introduction

Breast cancer is the most common cancer in women, with approximately 287,850 women in the United States alone being diagnosed in 2022 [1]. Increased awareness, early detection with frequent screenings, and expanded treatment options have improved breast cancer survival rates over time, with recent 5-year survival rates reported to be 90.6% [2]. Given these high survival rates, increased focus has been placed on long-term outcomes and patient quality of life after treatment. At present, the lowest rates of cancer recurrence are associated with surgical treatment options [3, 4]. As a result, breast cancer patients and their surgeons are often faced with choosing between breast-conserving surgery (BCS; otherwise known as lumpectomy) or mastectomy (removal of the whole breast), a decision-making process that is challenging, multi-faceted, and stressful. In recent years, BCS has replaced mastectomy as the preferred standard of care for early-stage breast cancer, since BCS has similar or improved survival rates and decreased risk of complications compared to mastectomy [5, 6, 7, 8]. With the goal of preserving healthy breast tissue and breast appearance, BCS involves the removal of the cancerous tissue along with a small margin of healthy tissue. As shown in Figure 1, the resulting tissue cavity undergoes a wound healing process that ultimately leads to variable levels of tissue contraction, scar tissue formation, and breast deformation (i.e., cosmetic defects, including dents, distortions, and asymmetries between breasts). The prognosis of a good cosmetic outcome typically weighs heavily on physician and patient selection of BCS over mastectomy, since good aesthetics has been associated with improved patient psychological recovery and quality of life [9, 10]. However, the complex nature of the tissue repair process as well as significant variations in patient-specific characteristics, make it extremely challenging, if not impossible, for surgeons to predict post-surgical healing, oncologic, and cosmetic outcomes. The inability to predict healing and breast deformation outcomes stems from the complex interplay between tissue mechanics, inflammatory-mediated biochemical and cellular signaling, and (myo)fibroblast mechanobiology during the tissue repair process. Therefore, there is a need for an improved mechanistic understanding of the multi-scale breast healing process along with definition of critical patient-specific characteristics that affect BCS outcomes. With this knowledge, surgeons and their patients can better develop individualized treatment plans that lead to decreased post-surgical complications, decreased surgical procedures (e.g., re-excision, revision, and/or reconstruction), and improved patient satisfaction and quality of life [5].

**Figure 1:**
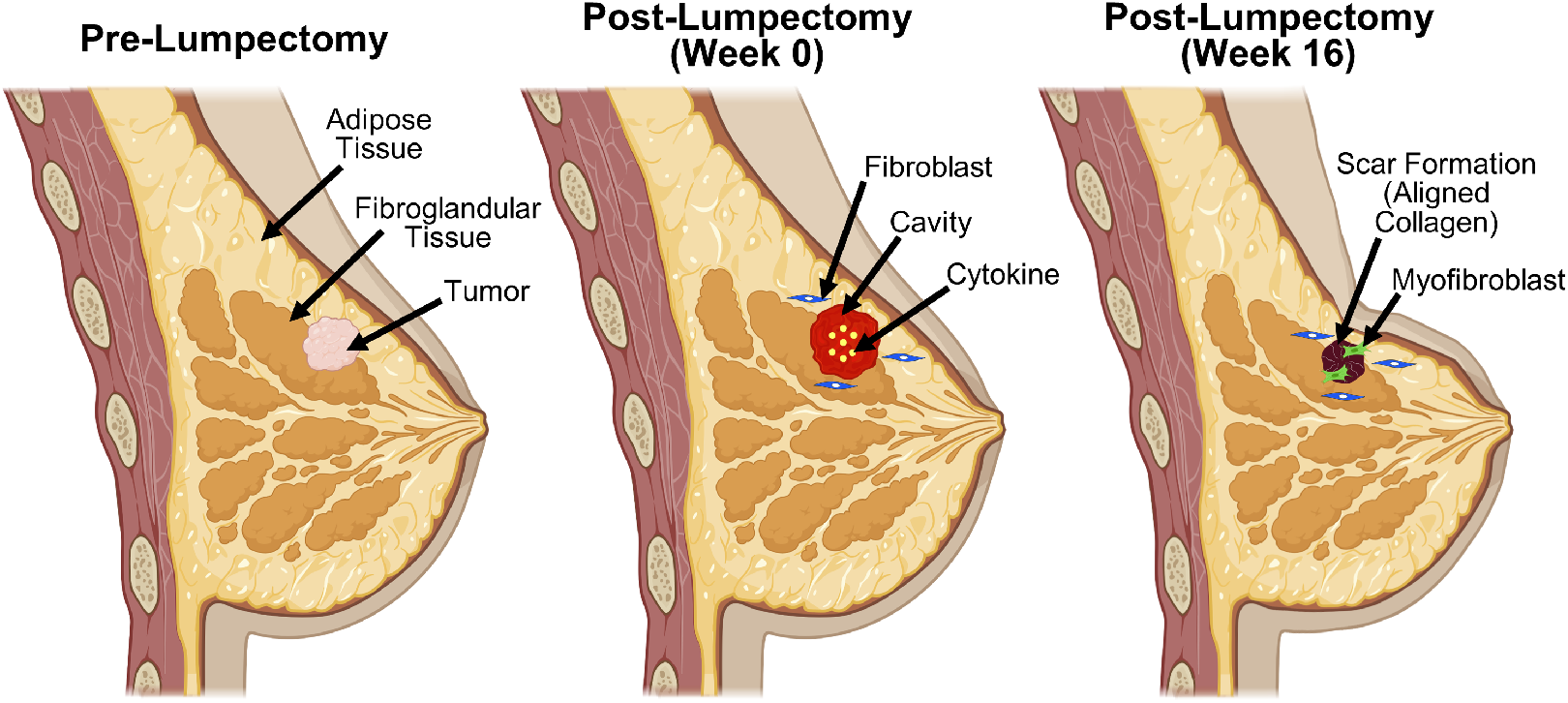
Schematic of cavity healing process following removal of breast tumor by lumpectomy. Tumor is excised along with a small margin of surrounding healthy tissue, forming a fluid-filled cavity. The surgical void undergoes the phased wound healing process, with hemostasis and inflammation phases resulting in creation of a cytokine gradient within the cavity. In turn, cytokines induce fibroblast migration and proliferation, resulting in collagen deposition and scar tissue formation through collagen fiber alignment. Fibroblast differentiation into myofibroblasts further promotes contraction of the cavity and surrounding tissue, which may contribute to breast deformities.

Given that few objective criteria and limited surgical decision-making tools exist, preoperative predictions of healing, oncologic, and breast cosmetic outcomes remain largely dependent on a surgeon’s past training and experience [11]. BCS surgical planning has been an evolving area over the past several years, as physicians work to further inform and standardize the process. In 2014 and 2016, the Society of Surgical Oncology (SSO), the American Society of Radiation Oncology (ASTRO), and the American Society of Clinical Oncology (ASCO), published consensus guidelines on adequate surgical margins when treating various types and stages of breast cancer with BCS and whole breast irradiation [12, 13]. Additionally, surgical decision trees have been developed based on correlative analyses of human BCS patient data, including tumor-to-breast volume percentage (TBVP), tumor location, breast cosmetic outcome assessments, and quality of life surveys [14, 15, 16]. While these decision-making tools provide recommendations on treatment thresholds (i.e., when to treat a patient with BCS versus mastectomy) based on tumor size and location, they have yet to receive widespread adoption. Feedback regarding patient satisfaction and quality of life, as provided through *BREAST* − *Q*^*TM*^ questionnaires and other patient surveys, has informed surgeons of other patient-specific factors affecting BCS outcomes [17]. More specifically, results from multivariable clinical analyses revealed that decreased breast density as measured by BI-RADS rankings, increased excised breast volume percentage -or equivalently cavity to breast volume percentage (CBVP)-, increased patient age, body mass index, breast irradiation, and concomitant adjuvant chemotherapy and radiotherapy often negatively influence surgical outcomes and patient satisfaction. [18, 19, 20, 21, 22]. In summary, since patient-specific characteristics are intertwined and significantly affect post-lumpectomy healing and cosmetic outcomes, there is a need for a predictive tool to better understand the mechanistic interplay between these contributing factors.

Computational models provide useful tools that can assist with informing, predicting, and simulating wound healing outcomes, including surgical wounds associated with BCS. In general, wound healing can be modeled as four, overlapping phases: hemostasis, inflammation, proliferation (or granulation), and remodeling [23]. To date, numerous numerical-based approaches have been developed to describe healing of superficial skin layers, including the epidermis and/or the dermis [24]. However, unlike skin wounds, which have an air-tissue interface, BCS yields a fully-enclosed cavity or void that resides relatively deep within the breast tissue. Healing of these deep, soft tissue wounds begins immediately following cavity creation, with blood clots (hematomas) and/or serous fluid (seromas) often filling the void [25]. The fibrin matrix, with its limited persistence and mechanical integrity, serves as a provisional scaffold, allowing local tissue contraction while promoting inflammation and cellularization. Platelet degranulation and cytokine secretion by inflammatory cells contributes to the formation of a cytokine gradient within the cavity, which, in turn, promotes fibroblast proliferation and migration into the wound space. Fibroblast proliferation, migration, and differentiation into myofibroblasts are further guided by fibrillar collagen deposition, and scaffold reorganization/contraction, ultimately creating a dense, stiff scar tissue within the contracted cavity. Scar tissue formation and remodeling over time are perhaps the most unpredictable aspects of BCS, since it is known to contribute to pain, breast deformations, and altered breast consistency, all of which negatively affect women emotionally and psychologically [26].

In recent years, computational models have also been developed for the purpose of predicting specific surgical outcomes following BCS. For example, Garbey and collaborators proposed a two-dimensional (2D) model to predict time-dependent changes in breast shape following lumpectomy [27, 28]. This model was calibrated using 1D MRI (magnetic resonance imaging) profiles obtained for a single patient [28]. Vavourakis and collaborators developed a 3D finite element model to predict breast deformation following BCS. Model validation was performed using a combination MRI and optical surface scans for 4 patients obtained before and 6 to 12 months after BCS [29]. Unfortunately, computational models developed to date lack a thorough calibration against experimental or clinical breast healing data. Additionally, present-day models do not fully capture the complex couplings between cellular mechanobiological activity, extracellular matrix (ECM) deposition and remodeling, and cavity and breast plastic deformation over time. Descriptions of collagen deposition, granulation tissue formation, and remodeling are especially important to capture, as the breast cavity and surrounding tissue will undergo large deformations and permanent contracture.

In this paper, we work to address this gap in wound mechanobiology modeling following BCS by presenting a theoretical and computational framework calibrated against animal model and clinical data. Here, we adapt our previously developed experimentally-calibrated model of dermal wounds that accounts for couplings between cellular mechanobiological activity, plastic deformations, and tissue remodeling [30, 31]. This informed 3D finite element model is then used to inform a machine learning surrogate model in order to evaluate the effect of specific mechanobiological parameters and patient-specific characteristics on healing and breast deformation outcomes. The proposed model has the potential to assist surgeons in creating an individualized treatment plan for patients that better predict oncologic, healing, and cosmetic outcomes.

## 2. Methods

The computational breast mechanobiological model represents a custom finite element solver implemented in C++. The link to the code repository is provided at the end of the manuscript. The software builds upon and extends our previous dermal wound healing models [23, 32, 30]. An overview of the model and associated adaptations is discussed below, with more detailed descriptions available in our previous work [32, 30]. Detailed parameter descriptions and values are included in Tables S1 and S2 in the Supplementary Material.

### 2.1. Geometry

We considered the two breast lumpectomy geometries shown in Figure 2. Both geometries were created and meshed in COMSOL (COMSOL Multiphysics, Burlington, MA). One geometry (Fig. 2A) corresponded to a generalized porcine breast based on a preclinical porcine lumpectomy study by Puls et al. (2021) [25]. Available ultrasound and explant images were used to estimate the dimensions of the ellipsoidal cavity (*a* = *b* = 1.5 cm, *c* = 0.6 cm) along with a cavity depth of 1.15 cm. The cavity represented approximately one-quarter of the total breast volume (quadrantectomy). The breast was assigned the shape of a half-ellipsoid (*a* = *b* = 2.32 cm, *c* = 2 cm), enclosed within a rectangular region (15 cm by 15 cm by 2 cm) of connective tissue.

**Figure 2:**
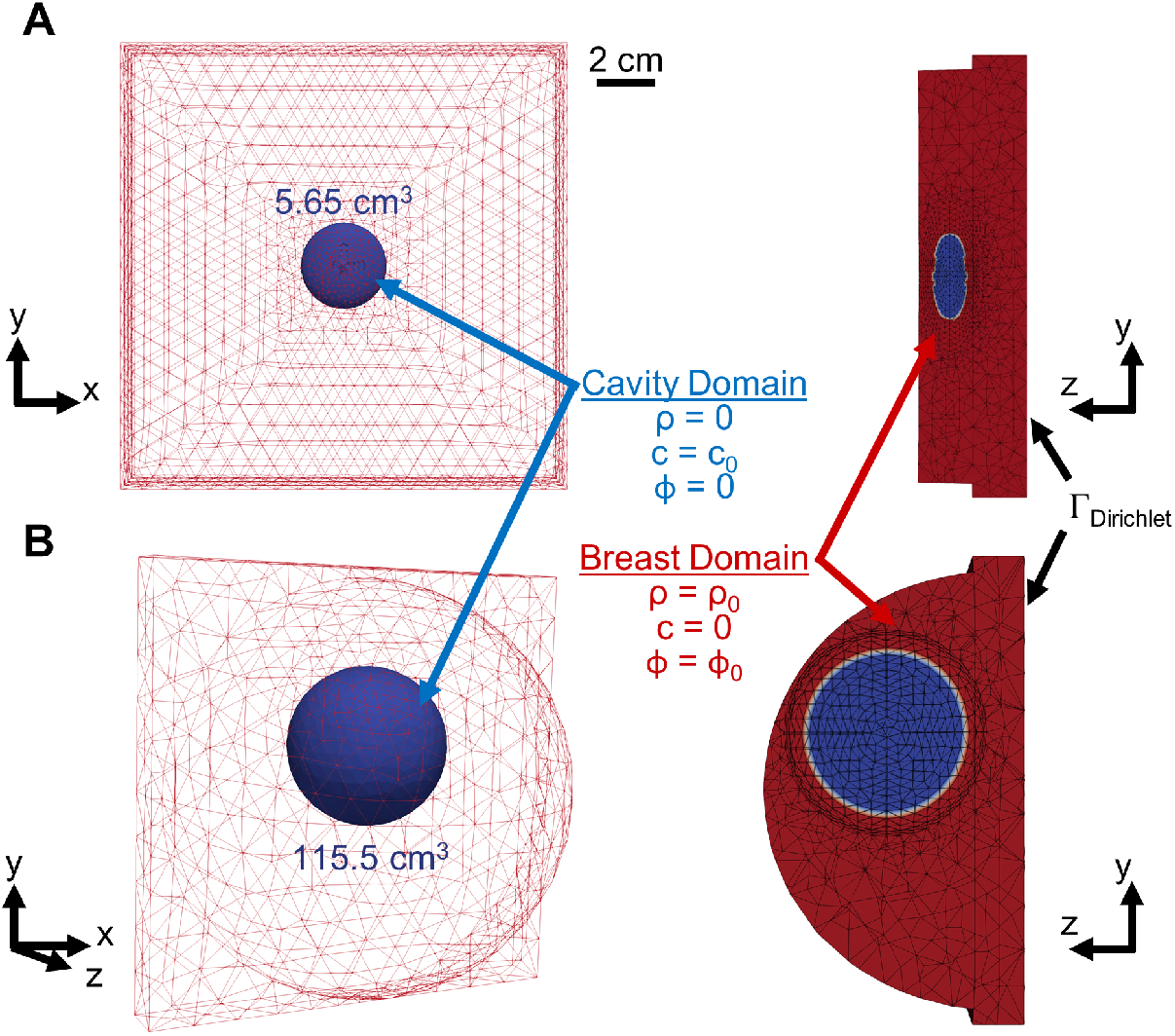
Meshing, initial conditions, and boundary conditions for the (A) porcine and (B) human breast geometries. For the porcine geometry, the breast was assumed to be a half-ellipsoid (22.60 cm^3^) and the cavity was assumed to be an ellipsoid (5.65 cm^3^), with both dimensions based on a quadrantectomy. The tissue external to the breast was modeled as connective tissue. For the human geometry, the breast was assumed to be a hemisphere with a volume of 1,324 cm^3^ and the cavity was assumed to be a sphere with a volume of 115.5 cm^3^. The Dirichlet boundary condition was applied to the interior surface of the 2-cm thick chest wall while the exterior surface of the breast was a free boundary.

An idealized human breast lumpectomy geometry was developed based on average breast and cavity sizes reported in a human clinical study by Prendergast et al. (2009) [33]. As shown in Figure 2B, the breast was modeled as a hemisphere with a radius of 8.58 cm and the cavity was modeled as a sphere with a radius of 3.02 cm. Since the upper outer quadrant is reported to be the most prevalent tumor location [34, 35, 33, 36, 15], this cavity location was assumed in the model. Breast cavity contraction over a four-week period following BCS, as quantified by Prendergast and co-workers, was also used for model calibration.

### 2.2. Kinematics

The reference geometries displayed in Figure 2 are described with material coordinates **X** ∈ ℬ_0_ ⊂ ℝ^3^. Through the deformation mapping *φ*, the time-dependent configuration, ℬ_*t*_, is obtained as **x** = *φ*(**X**, *t*). The fibroblast density, cytokine concentration, and collagen density are *ρ*(**x**, *t*), *c*(**x**, *t*), *ϕ*(**x**, *t*), respectively. The collagen matrix is further defined through the fiber dispersion *κ*(**x**, *t*) and the preferred fiber orientation **a**_0_(**x**, *t*). The deformation gradient **F** = *∂***x***/∂***X**, which describes local geometry changes, can be split into two separate components capturing the elastic and plastic deformation

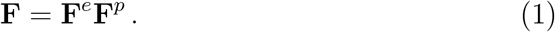

Furthermore, the plastic deformation tensor is described with three scalar fields

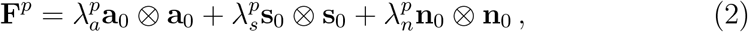

where vectors **a**_0_, **s**_0_, **n**_0_ form an orthonormal basis around the preferred fiber orientation **a**_0_.

### 2.3. Constitutive and balance equations

The change in the fields introduced in the previous section are classified into three categories. The *biological* fields *ρ, c* satisfy mass balance in the form of reaction-diffusion partial differential equations (PDEs). The *microstructural* fields *ϕ*, 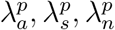, *κ*, **a**_0_ do not have a diffusion component and their change is local. The microstructural fields are directly coupled to the *mechanical* field of deformation *φ*, which satisfies momentum balance.

#### 2.3.1. Biochemical model

Fibroblast density and cytokine concentration satisfy standard advectiondiffusion transport equations

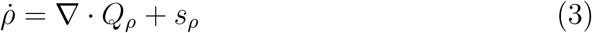

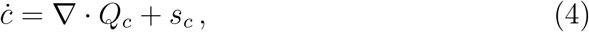

where *Q*_*ρ*_, *Q*_*c*_ are flux terms akin to Fickian diffusion

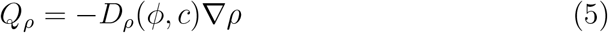

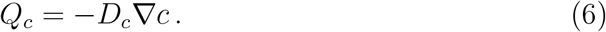

While the diffusion coefficient for the cytokine is assumed constant, cell diffusion (migration) is affected by both cytokine concentration and collagen densities,

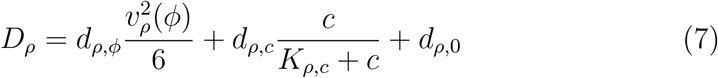

with parameters *d*_*ρ,ϕ*_, *d*_*ρ,c*_, *d*_*ρ*,0_. The first term in eq.(7) reflects the direct dependence of fibroblast speed on collagen density, while the second and third terms are related to the baseline diffusion coefficient for cells in native tissue and their change in diffusivity with *c* considering Michaelis Menten kinetics. The initial profile for *v*_*ρ*_(*ϕ*) was estimated through available in-vivo wound healing data [25, 37]. The expression was then modified through a parameter Δ, which skews the collagen concentration associated with maximum fibroblast speed. Additional information about *v*_*ρ*_(*ϕ*) and Δ can be found in Figure S1 in the Supplementary Material.

The source terms *s*_*ρ*_, *s*_*c*_ are

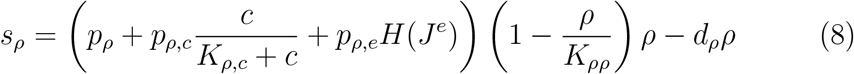

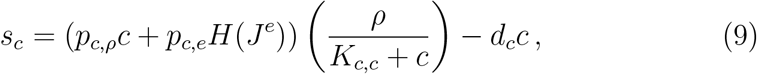

with parameters *p*_*ρ*_, *p*_*ρ,c*_, *K*_*ρ,c*_, *p*_*ρ,e*_, *K*_*ρρ*_, *d*_*ρ*_ for the fibroblast source and *p*_*c,ρ*_, *p*_*c,e*_, *K*_*c,c*_, *d*_*c*_ for the cytokine. The values of all parameters are listed in Table S1 in the Supplementary Material.

Note that most dependencies of the biological fields are on other biological fields, but some couplings exist in the microstructural and mechanical fields. For instance, cell migration in eq. (7) depends on the microstructural field *ϕ* through *v*_*ρ*_ defined in the Supplementary Material. The biological fields are also coupled to the mechanical field through the mechanosensing logistic function, *H*(*J*^*e*^) in eqs. (8) and (9) described below.

#### 2.3.2. Mechanical model

Balance of linear momentum in the absence of body force is reduced to the standard equation

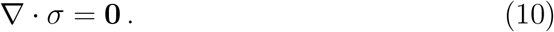

However, here the total stress is split into two separate components for active and passive stress contributions

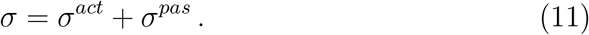

The active stress is described in the following section devoted to the mechanobiological couplings. In this section, we focus on the passive part. The passive material response is assumed hyperelastic with the strain energy function

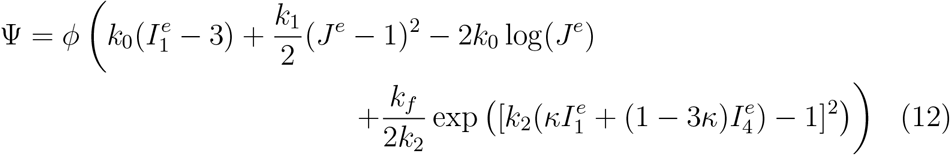

parameterized by *k*_0_, *k*_1_, *k*_2_, *k*_*f*_. It is also a function of the microstructure fields *ϕ, κ*, and of the elastic invariants of the deformation 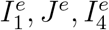. Note that only the elastic part of the deformation contributes to the strain energy. Based on the split eq.(1), the elastic volume change is *J*^*e*^ = det(**F**^*e*^), the first isotropic invariant is the trace of the elastic right Cauchy Green tensor 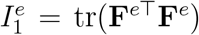, and the fourth invariant describes the deformation in the preferred fiber direction 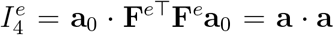, with **a** representing the deformed fiber orientation.

The parameters *k*_0_ and *k*_1_, which correspond to a neo-Hookean contribution, were determined using the rule of mixtures assuming that human and porcine breast tissue, on average, is composed of 70% adipose tissue and 30% fibroglandular tissue [38]. Han et al. (2011) and several other studies were used to inform material properties for adipose and fibroglandular tissue, as we estimated Young’s modulus for adipose and fibroglandular tissue to be 10 kPa and 40 kPa, respectively [39, 40, 41, 42, 43, 44, 45, 46, 47]. The parameter *k*_*f*_ denotes collagen fiber stiffness for the scar tissue [48]. Mechanical parameter descriptions and values are included in Table S2 in the Supplementary Material.

#### 2.3.3. Mechanobiological coupling

As mentioned before, the biological fields are linked to the mechanical deformation by the logistic function *H*(*J*^*e*^) in eqs. (8) and (9). This function encodes a mechanosensing activation as the deformation deviates from homeostasis

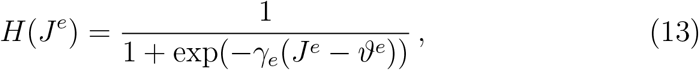

with parameters *γ*_*e*_, *ϑ*^*e*^. Another coupling that appeared in eq. (11) is the active stress, which is defined as

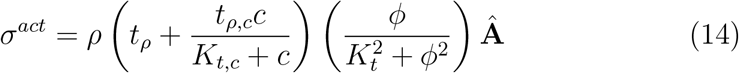

which depends on the fibroblast density *ρ*, the cytokine *c*, the collagen density *ϕ*, and the preferred fiber orientation through the structure tensor **Â** = **A***/*tr(**A**), **A** = **I** + (1 − 3*κ*)**a** ⊗ **a**. The parameters of the active stress eq. (14) are *t*_*ρ*_, *t*_*ρ,c*_, *K*_*t*_, *K*_*t,c*_, with parameter descriptions and values provided in Table S2 in the Supplementary Material.

The other mechanobiological coupling that was introduced earlier is the fibroblast migration dependence on collagen density in a non-monotonic fashion through *v*_*ρ*_ in eq. (7) [30].

The last set of equations needed to close the model are the rate equations for the microstructural fields. Collagen deposition is encoded by

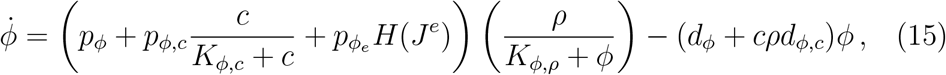

with dependence on both cell density and cytokine concentration. Descriptions and values of parameters 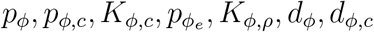) in eq. (15) are included in Table S2 in the Supplementary Material. The change in plastic deformation occurs independently in all three directions

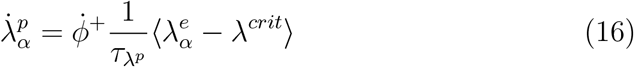

where *α* = {*a, s, n*} are the three directions of the orthonormal frame **a**_0_, **s**_0_, **n**_0_. The term 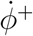 in eq. (16) is the positive part of the rate of change of collagen (i.e., the new collagen deposition rate), which contributes to deformation plastification. The Macaulay brackets ⟨•⟩ specify that plastic deformation only occurs beyond some threshold deformation *λ*^*crit*^.

Lastly, the change in preferred collagen fiber orientation and dispersion are based on the eigenvalues of the deformation

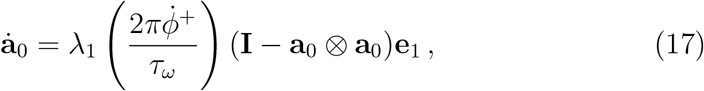

where *λ*_1_, **e**_1_ are the largest eigenvalue and corresponding eigenvector, respectively. Eq. (17) essentially reorients the principal fiber direction to the direction of maximum principal stretch, with time constant *τ*_*ω*_ dependent on collagen deposition 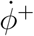. The fiber dispersion change

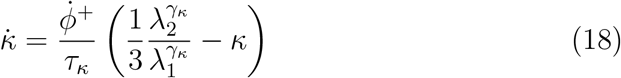

depends on the ratio of the first two eigenvalues with a power law parameterized by *γ*_*κ*_ and the time constant *τ*_*κ*_.

### 2.4. Experimental data

Time-dependent changes in fibroblast and collagen densities were informed by histopathological data from the porcine lumpectomy study [25]. Hematoxylin and eosin (H&E) stained cross-sections of breast explants were analyzed 1 week, 4 weeks, and 16 weeks following lumpectomy and compared to normal porcine breast tissue (Fig. 3). An image of each cross-section was post-processed in Aperio ImageScope (Leica Biosystems, Vista, CA) and 25 individual regions (500 × 500 *µ*m^2^) spanning the cavity domain were extracted. These regions were further processed in ImageJ (National Institutes of Health, Bethesda, MD), where multiple color balance filters were applied to quantify the number fibroblasts, red blood cells (RBCs), and immune cells per region. Fibroblast number per area was used to calculate fibroblast volume density, assuming a tissue section thickness of 4 *µ*m. Additional details of this image analysis process are provided in the Supplementary Material. The H&E stained cross-sections were also used to determine collagen density by correlating collagen density with the intensity of eosin-stained collagen fibers. Eosin intensity for a region of interest was determined using ImageJ and normalized to connective tissue values within adjacent healthy breast tissue values. When calculating normalized collagen densities, an average breast composition of 70% adipose tissue and 30% fibroglandular tissue was assumed [38].

**Figure 3:**
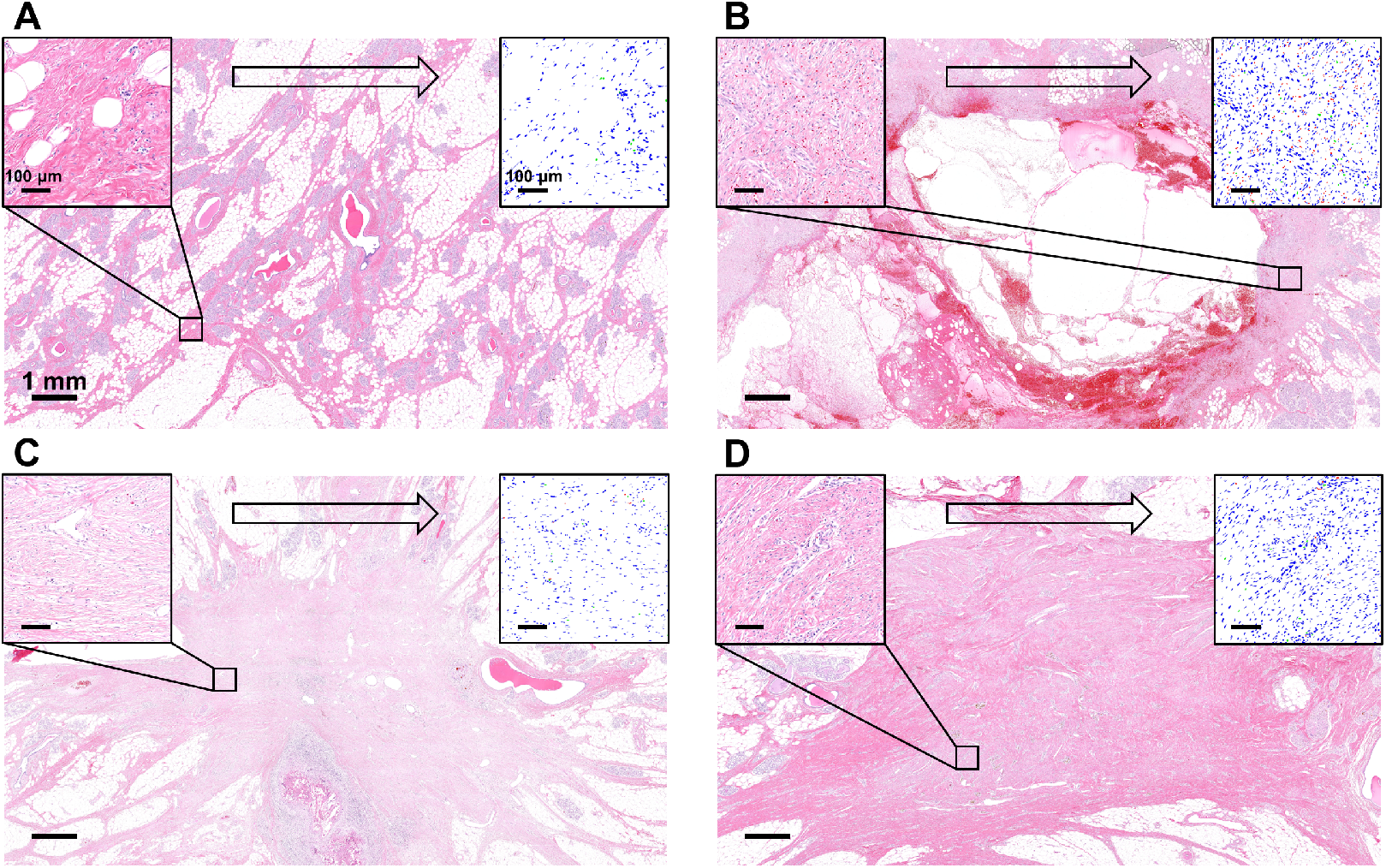
Overview of histological image analysis process used to quantify fibroblast and collagen densities within (A) normal porcine breast tissue and porcine breast tissue undergoing progressive healing at (B) 1 week, (C) 4 weeks, and (D) 16 weeks following simulated lumpectomy (quadrantectomy). Individual regions (500 × 500 *µm*^2^) of H&Estained cross-sections (top left inset) were processed using a particle analyzer (top right inset) for identification and enumeration of fibroblasts (blue), RBCs (red), and immune cells (green). Collagen density was determined by normalizing regional eosin intensity values for connective tissue within healing breasts to eosin intensity in normal breast connective tissue.

Temporal changes in cytokine concentration were informed by prior human clinical studies that evaluated cytokine levels in seroma fluid, which commonly fills the breast void following surgery. Seroma fluid is known to be composed of cytokines that impact the inflammation and proliferation phases of healing [49]. It has also been reported that seromas formed following BCS resolve within approximately 4 weeks [50]. Based on this, it was assumed that cytokine levels decayed exponentially over approximately a 4-week time period.

### 2.5. Model calibration using Gaussian processes

The finite element model defined in previous sections is computationally expensive and impractical for tasks such as model calibration or sensitivity analysis. Therefore, to calibrate the model against experimental porcine data and human clinical data, we leveraged Gaussian process (GP) surrogates [51]. The methodology for GP model calibration is illustrated in Figure 4. Calibration was performed with two separate GPs. First, a submodel consisting only of the biological fields *ρ, c* and the microstructural field *ϕ* was isolated out of the complete set of equations with the goal of fitting the porcine histology data (i.e., fibroblast and collagen densities). A second GP was constructed for the fully coupled mechanobiological model. This two-stage approach was used to i) inform biological parameters that could, in turn, be compared with other computational models lacking mechanobiological couplings, and ii) calibrate the mechanobiological coupling terms, for which limited prior information exists.

**Figure 4:**
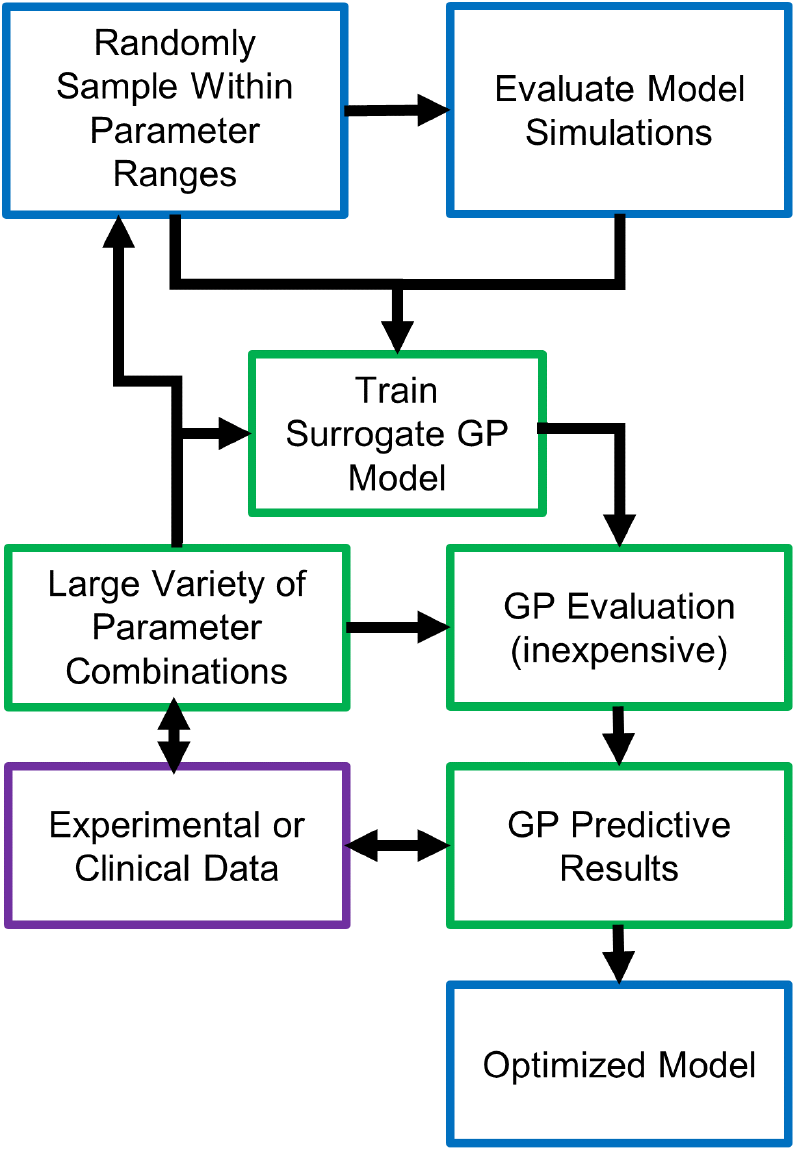
GP methodology used to identify optimum biochemical and mechanobiological parameters that best fit porcine lumpectomy histology results and human clinical contraction data. The computational model was run several times, sampling across the entire parameter space to train the GP model. The GP model was then used minimize RMSE by comparing GP generated curves against experimental and clinical data. Regions of interest within the parameter space (because they were predicted to minimize RMSE and had a large predicted variance) were further sampled, finite element simulations run, and GP model updated for further minimization.

For the first GP surrogate, 5 parameters Θ_*b*_ = {*p*_*ρ,c*_, *d*_*ρ,ϕ*_, Δ, *p*_*ϕ*_, *p*_*ϕ,c*_} were sampled from the ranges reported in Table 1 using Latin Hypercube Sampling (LHS). All other parameters affecting the submodel {*ρ, c, ϕ*} were assigned values from literature or calculated in order to satisfy a physiological steady state. In other words, the 5 parameters Θ_*b*_ were identified as the adjustable parameters for model calibration. To train the GP, 100 different parameter combinations of Θ_*b*_ were generated and applied to the finite element submodel, with fibroblast and collagen density values at the center of the cavity *ρ*_*C*_(**t**), *ϕ*_*C*_(**t**) representing model outcomes of interest. A total of 196 time steps were extracted from the simulation, covering the time *t* ∈ [0, 16] weeks. Following calibration, the GP model was used for minimization of root mean square error (RMSE) by comparing GP predictions for 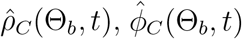 against porcine histopathological data. After minimization, regions of the parameter space Θ_*b*_ with lower RMSE and higher predicted variance were used to select new Θ_*b*_ parameter combinations to further train the GP model. Subsequent RMSE minimization with the GP model yielded the optimal parameter values Θ_*b*_.

**Table 1:** Biochemical and mechanobiological parameters with established initial ranges that were evaluated and optimized using the biochemical or mechanobiological GP.

| Parameter | Description | Range | Optimized Value |
| --- | --- | --- | --- |
| **Biochemical GP Parameters ($\Theta_b$)** |  |  |  |
| $p_{\rho,c}$ [1/hr] | Cytokine-Increased Proliferation | [0.0092641, 0.04632] | 0.015314 |
| $d_{\rho,\phi}$ [-] | Fibroblast Diffusion Scaling Constant | [62.6793, 12472.9067] | 1582.3 |
| $\Delta$ [-] | Skewness of Fibroblast Speed $v_\rho(\phi)$ | [0, 1] | 0 |
| $p_\phi$ [1/hr] | Collagen Production | $[3.633 \times 10^{-9}, 3.633 \times 10^{-7}]$ | $1.4 \times 10^{-8}$ |
| $p_{\phi,c}$ [1/hr] | Collagen Production Activated by Cytokine | $[3.633 \times 10^{-9}, 3.633 \times 10^{-7}]$ | $7.0 \times 10^{-8}$ |
| **Mechanobiological GP Parameters ($\Theta_m$)** |  |  |  |
| $t_\rho$ [MPa] | Contractile Force of Fibroblasts | $[9.08244 \times 10^{-8}, 5.44947 \times 10^{-7}]$ | $2.33548 \times 10^{-7}$ |
| $t_{\rho,c}$ [MPa] | Contractile Force of Myofibroblasts | $[1 \cdot t_\rho, 5 \cdot t_\rho]$ | $3.28571 \cdot t_\rho$ |
| $K_t$ [-] | Saturation of Mechanical Force by Collagen | [0.1, 0.5] | 0.2 |
| $\tau_{\lambda^p}$ [1/hr] | Rate of Plastic Deformation | [0.00485, 0.2425] | 0.05 |

After calibration of the {*ρ, c, ϕ*}-submodel, a similar approach was performed to calibrate the mechanobiological parameters Θ_*m*_ = {*t*_*ρ*_, *t*_*ρ,c*_, *K*_*t*_, *τ*_*λp*_}. For the second GP model, a total of 100 simulations were run after LHS sampling of Θ_*m*_ within the specified ranges in Table 1. The trained GP was used to minimize the RMSE with respect to the cavity contraction data from the human clinical study [33]. As described previously, initial minimization was followed by subsequent finite element model parameter evaluations and training of the GP model.

## 3. Results

### 3.1. Pathophysiologic findings through porcine histology analysis

Analysis of breast histological cross-sections from a longitudinal porcine lumpectomy study informed fibroblast and collagen densities within the breast cavity at 1, 4, and 16 weeks after surgery. Table 2 summarizes values for each post-surgical time point compared to healthy breast tissue. Given that hematomas or seromas were observed grossly and histologically 1 week following lumpectomy (Fig. 3B), fibroblast and collagen densities were assumed to be zero for this time point. By 4 weeks, fibrovascular scar tissue was evident within the contracted cavity (Fig. 3C), with fibroblast and collagen density values roughly 7 and 1.3 times healthy breast tissue values, respectively. By 16 weeks, the fibrous scar tissue increased in collagen density (approximately 2.3 times healthy breast tissue values), appearing as differentially oriented swirls of parallel-aligned fibers (Fig. 3D). Although fibroblast density decreased between 4 and 16 week time points, values remained high at roughly 4 times those for healthy breast tissue.

**Table 2:** Fibroblast and collagen densities (mean ± SD) quantified from histological crosssections of normal, healthy porcine breast tissue and explanted breast tissue at 1 week, 4 weeks, and 16 weeks following lumpectomy. Post-surgical values represent the cavity center.

| Time Point | Fibroblast Density<br>(Mean ± SD) [$cells/mm^3$] | Collagen Density<br>(Mean ± SD) [$\phi/\phi_0$] |
| --- | --- | --- |
| Healthy Tissue | 55,051 ± 15,527 | 1 ± 0 |
| 1 Week Post-Surgery | 0 ± 0 | 0 ± 0 |
| 4 Weeks Post-Surgery | 377,504 ± 94,279 | 1.35 ± 0.25 |
| 16 Weeks Post-Surgery | 215,893 ± 45,150 | 2.33 ± 0.35 |

### 3.2. Calibration of the {ρ, c, ϕ} submodel

Fibroblast and collagen density values reported in Table 2 were successfully fit to the {*ρ, c, ϕ*} submodel by optimizing the (Θ_*b*_). Predicted fibroblast and collagen density values fell within experimentally-determined standard deviation ranges for all time points (Fig. 5). Finite element simulations for the optimized submodel are shown in Figure 5, illustrating spatiotemporal changes in fibroblast density, collagen density, and cytokine concentration.

**Figure 5:**
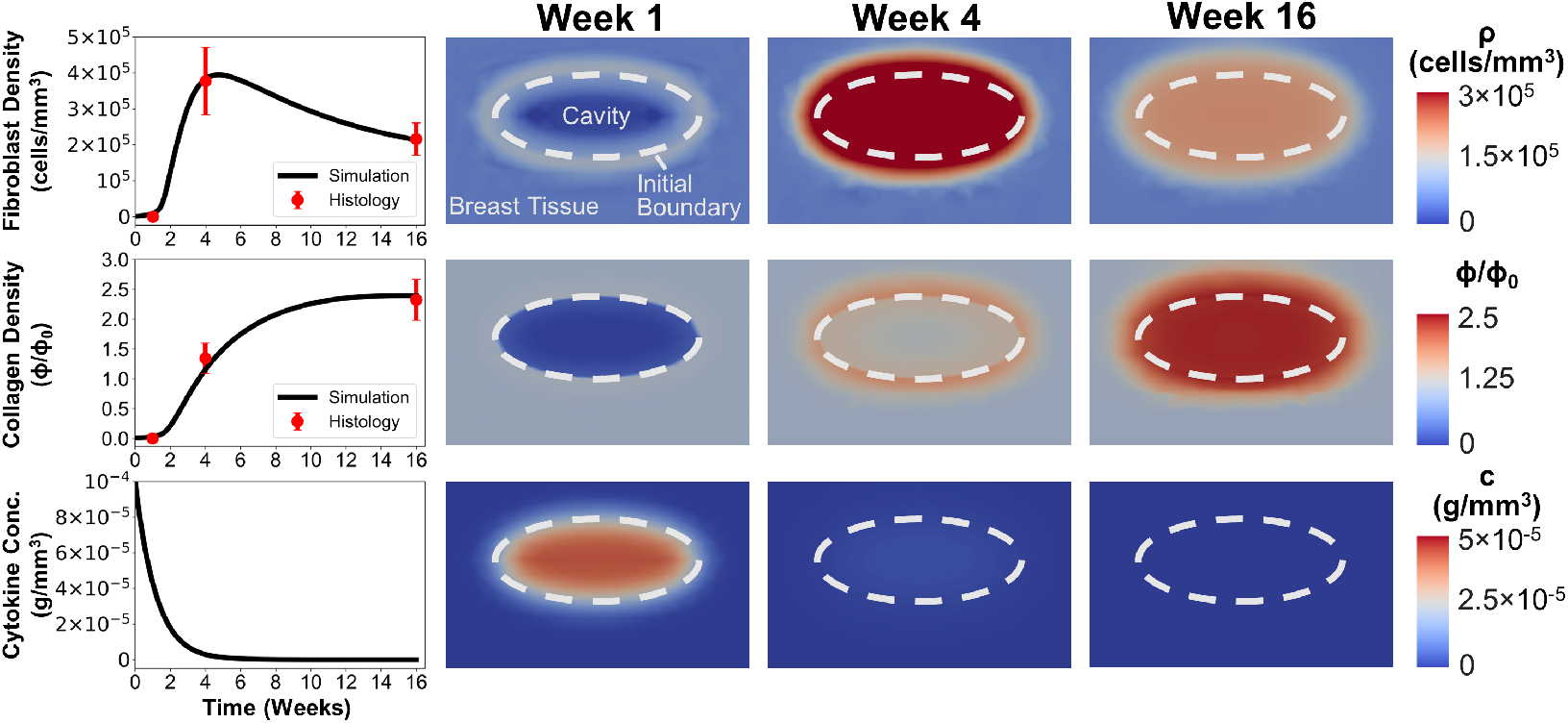
Simulation results of the {*ρ, c, ϕ*} submodel using optimized parameters Θ_*B*_. Plots display time-dependent changes in fibroblast density, collagen density, and cytokine concentration at the cavity center as determined from simulations and histology. Corresponding contour plots from breast cavity healing simulations are shown for weeks 1, 4, and 16.

Fibroblast and collagen densities within the cavity center were roughly zero at week 1 of the simulation (Fig. 5), successfully modeling hematoma and/or seroma formation and the lack of fibroblast infiltration observed histologically (Fig. 3B). Contour plots showed modest increases in fibroblast and collagen density, respectively, at the cavity-tissue interface (Fig. 5), which also matched histological findings (Fig. 3B). Fibroblast density increased sharply between weeks 1 and 4 (Fig. 5), effectively simulating fibroblast proliferation and migration. An increase in collagen density followed thereafter (Fig. 5), which is consistent with progressive collagen deposition by fibroblasts during the proliferation phase of healing. As shown in Figure 5, simulation results reached a maximum fibroblast density of 3.95 × 10^5^ *cells/mm*^3^ at roughly 4.5 weeks, after which time fibroblast density steadily declined to match histological outcomes. As fibroblast number declined between 4 and 16 weeks, the rate of collagen deposition declined, with collagen density values plateauing within experimentally measured ranges (Fig. 5). Simulated cytokine concentration within the cavity started at the maximum nominal value and showed a rapid decay over the first four weeks (Fig. 5). Such results are consistent with events and phases of wound healing as reported in the literature [52, 50].

### 3.3. Calibration of the fully coupled mechanobiological model

Human breast cavity contraction data estimated from Prendergast et al. (2009) was fit with the coupled mechanobiological model by optimizing mechanobiological parameters (Θ_*m*_) listed in Table 1. Results from the calibrated finite element simulation, including cavity contraction, permanent deformation, and breast surface deformation, are displayed in Figure 6. Consistent with human data, the simulated post-surgical breast cavity contracted to approximately 66.49% of its original cavity volume within 1 week. The cavity volume continued to decrease, contracting to 20.90% of its original volume in just 16 days following surgery. By 4 weeks, the cavity showed a modest increase in volume to reach 31.43% of the excised volume. The overall shape of the contraction curve was similar to porcine lumpectomy study findings as well as cavity contraction in human patients following BCS and whole-breast irradiation [25, 53].

**Figure 6:**
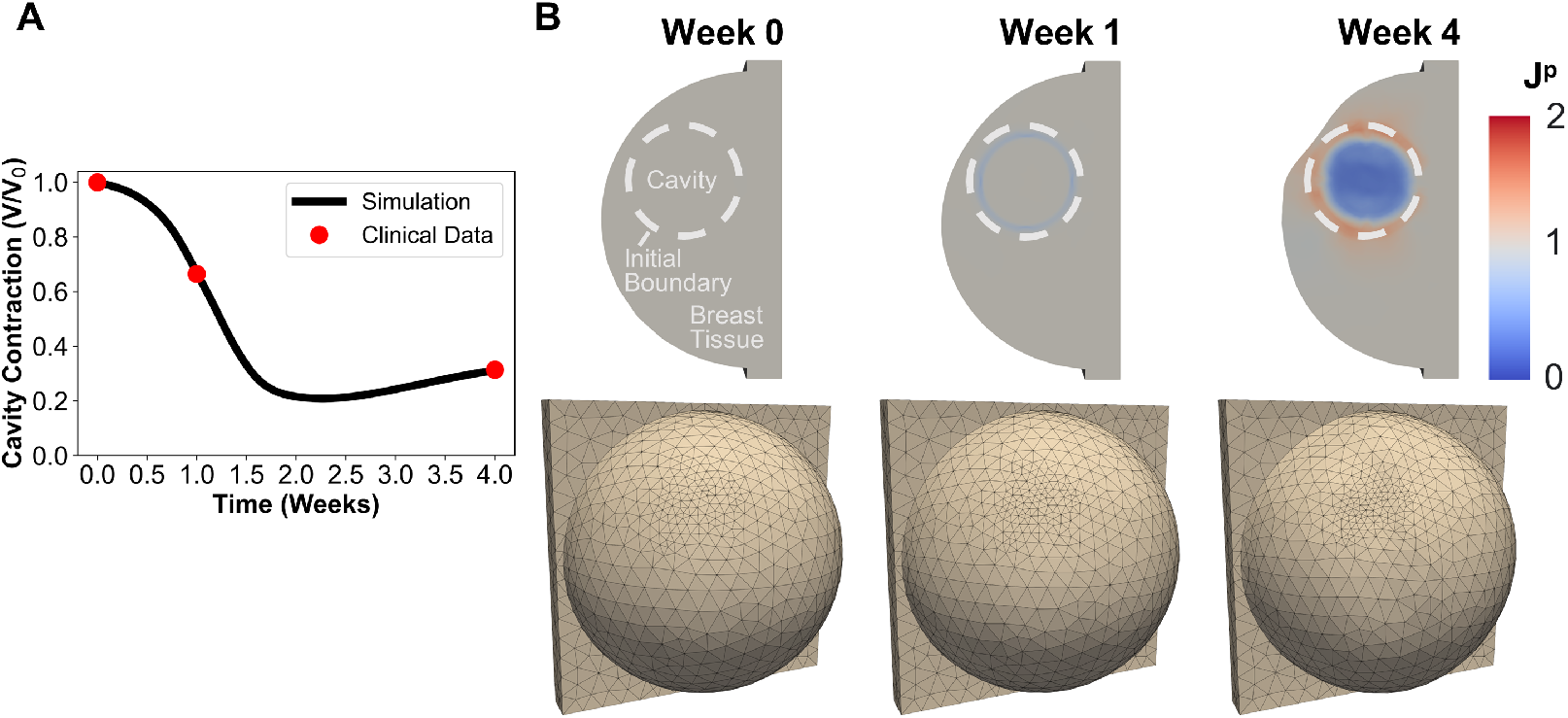
Mechanobiological model outcomes using optimized parameters Θ_*m*_. (A) Simulated post-surgical cavity contraction over time compared to clinical data. (B) Contour plots displaying time-dependent changes in permanent tissue deformation for simulated breast cavity healing (top) and associated breast surface deformation (bottom).

Permanent deformation (*J*^*p*^) was also visible across the cavity domain and surrounding tissue, leading to breast surface deformations (Fig. 6B). At the time of tumor removal (t = 0 week), no change in tissue volume is observed across the entire geometry (*J*^*p*^ = 1). Immediately thereafter, permanent contracture (*J*^*p*^ *<* 1) becomes prevalent at the tissue-cavity interface, with *J*^*p*^ = 0.85 for this region at the 1-week time point. This permanent deformation contributed to a modest surface asymmetry in the upper outer quadrant breast (Fig. 6B). By week 4, severe permanent contracture (*J*^*p*^ = 0.3) was observed within the cavity while tissue surrounding the cavity was experiencing tensional forces (*J*^*p*^ *>* 1) directed perpendicular to the cavity surface. Such observations are consistent with tissue repair and scar formation, as newly deposited collagen fibers within the cavity are contracted and reoriented by fibroblasts and myofibroblasts and the surrounding tissue ECM is drawn in tension [30, 31]. This permanent contracture contributed to an obvious breast surface deformity adjacent to the cavity (Fig. 6B).

### 3.4. Mechanobiological parameter sensitivity analysis

A major goal associated with the calibration of our detailed mechanistic model of breast healing after BCS is to better define key parameters and relationships that influence healing and cosmetic outcomes. In particular, mechanobiological model calibration, as described in previous sections, allowed optimization of parameters Θ_*m*_ for which there is little direct experimental or clinical information. An important next step was to explore the sensitivity of model predictions with respect to these parameters. To analyze Θ_*m*_ parameter effects, 2500 predictive cavity contraction curves were generated with the calibrated GP by sampling Θ_*m*_ values within ranges reported in Table 1. The normalized cavity volume at week 4 (*V*_4_*/V*_0_) was probed, with Figure 7A-D showing four 2D contour plots where the force of fibroblasts (*t*_*ρ*_), force of myofibroblasts (*t*_*ρ,c*_), saturation of mechanical force by collagen (*K*_*t*_), and rate of plastic deformation (*τ*_*λp*_) were varied.

**Figure 7:**
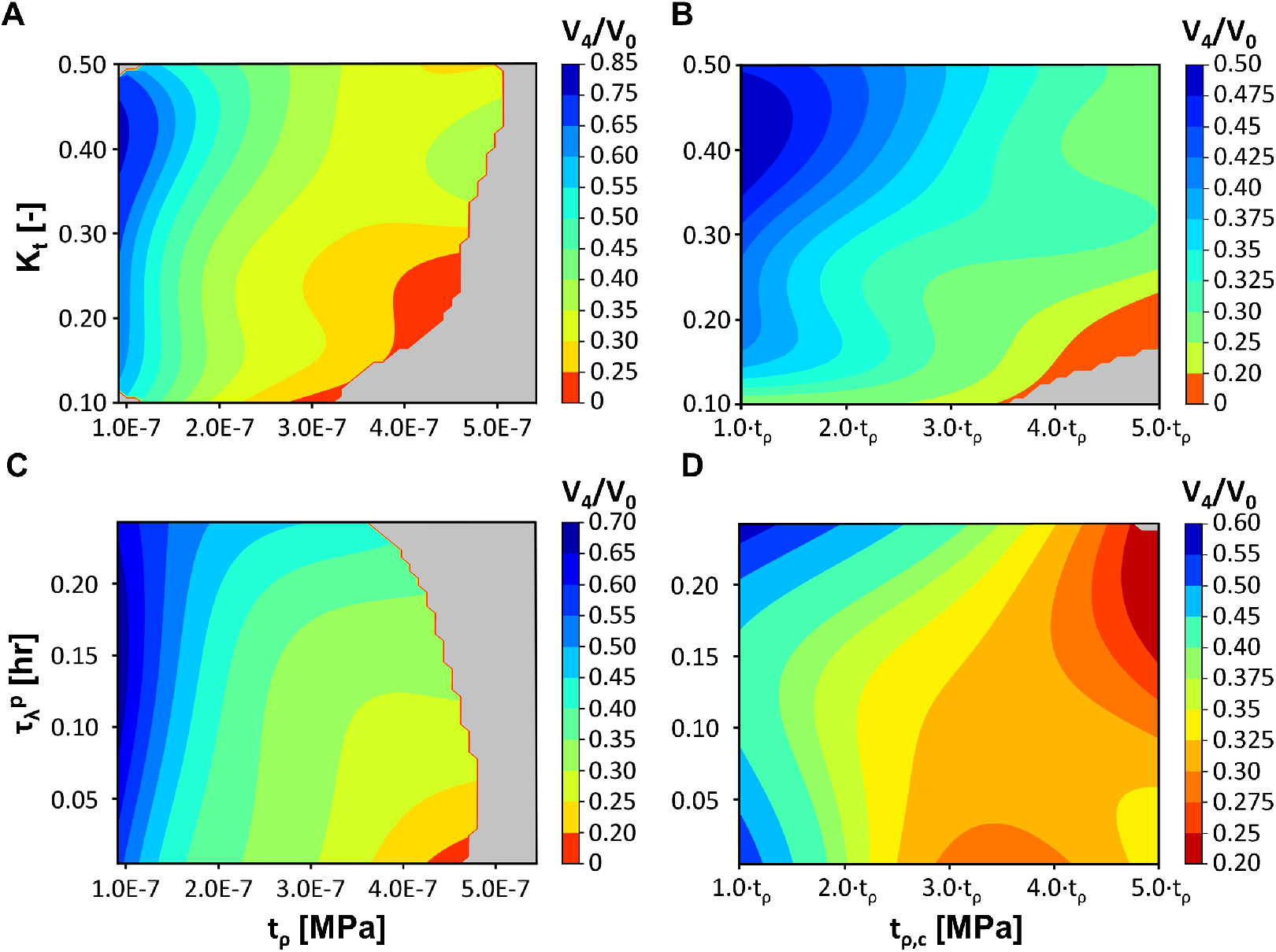
Plots showing relationships between mechanobiological parameters *t*_*ρ*_, *t*_*ρ,c*_, *K*_*t*_, and *τ*_*λp*_. Plots were created based on predictive cavity contraction curves generated using the mechanobiological GP model by varying two of the four parameters (constants used: *t*_*ρ*_= 1.5 × 10^*−*5^ MPa, *t*_*ρ,c*_ = 2.5 · *t*_*ρ*_ MPa, *K*_*t*_= 0.3, and *τ*_*λp*_ = 0.1 hr) and evaluating the change in cavity volume at week 4. Gray regions on the plots represent regions in the parameter space that were not well informed and for which the predicted variance by the GP model were large.

As shown in Figure 7A, cavity contraction was highly dependent on the fibroblast force *t*_*ρ*_, with increasing force leading to larger contraction. Although *K*_*t*_ had a less pronounced effect, increasing the saturation of mechanical force by collagen was found to decrease cavity contraction. Due to this inverse relationship, low *K*_*t*_ values and high *t*_*ρ*_ values produced the largest contractions, with the cavities contracting to less than 25% of their initial volume by week 4. Cavity contraction also increased with increasing myofibroblast force *t*_*ρ,c*_; however, an interesting coupling was identified between *K*_*t*_ and *t*_*ρ,c*_ (Fig. 7B). Evaluation of *t*_*ρ*_ and *K*_*t*_ pairings (Fig. 7A) clearly showed that fibroblast force was the dominant parameter. By contrast, results for *t*_*ρ,c*_ and *K*_*t*_ pairings (Fig. 7B) suggested that collagen saturation (*K*_*t*_) had a more pronounced effect coupled to *t*_*ρ,c*_ at lower *K*_*t*_ values. For example, when *K*_*t*_ = 0.1, cavity contraction values ranged between 25% and 30% for *t*_*ρ,c*_ ∈ [1 · *t*_*ρ*_, 2.5 · *t*_*ρ*_]. A broader cavity contraction range was observed for *K*_*t*_ = 0.5, with values varying from 50% to 37.5% across *t*_*ρ,c*_ ∈ [1 · *t*_*ρ*_, 2.5 · *t*_*ρ*_].

The rate of plastic deformation (*τ*_*λp*_) was inversely related to cavity contraction. In other words, lower values of *τ*_*λp*_ supported larger cavity contraction. The contour plot showing *τ*_*λp*_ and *t*_*ρ*_ pairings (Fig. 7C) revealed that cavity contraction was less sensitive to *τ*_*λp*_ for lower *t*_*ρ*_ values. However, as *t*_*ρ*_ increased, the rate of plastic deformation became more influential on contraction outcomes. For the *τ*_*λp*_ versus *t*_*ρ,c*_ contour (Fig. 7D), it was found that myofibroblast force was tightly coupled to the rate of plastic deformation, with cavity contraction becoming more severe for lower *τ*_*λp*_ and larger *t*_*ρ,c*_ values. Interestingly, the greatest cavity contraction (between 20% to 25%) occurred when both *τ*_*λp*_ and *t*_*ρ,c*_ had larger values.

### 3.5. Effect of cavity-to-breast volume percentage

Since the mechanobiological model was informed based on human BCS cavity contraction data, it can be applied to predict how patient-to-patient variability in breast and tumor characteristics affect healing and cosmetic outcomes. For example, the effect of CBVP was evaluated to identify trends in spatiotemporal cavity contraction and breast deformation. This model application involved adding CBVP as an input variable to the established mechanobiological GP. Similar to the initial GP model calibration, LHS sampling of the parameters Θ_*m*_ and CBVP was performed. Following GP model re-calibration, 2,500 GP predictive contraction curves were then used to evaluate the 4-week post-surgical cavity contraction and breast deformation for CBVP values between 0.43% and 8.7% (Fig. 8A). This CBVP range was based on geometric constraints of the assumed breast geometry and captures the wide range of reported breast tumor sizes [54].

**Figure 8:**
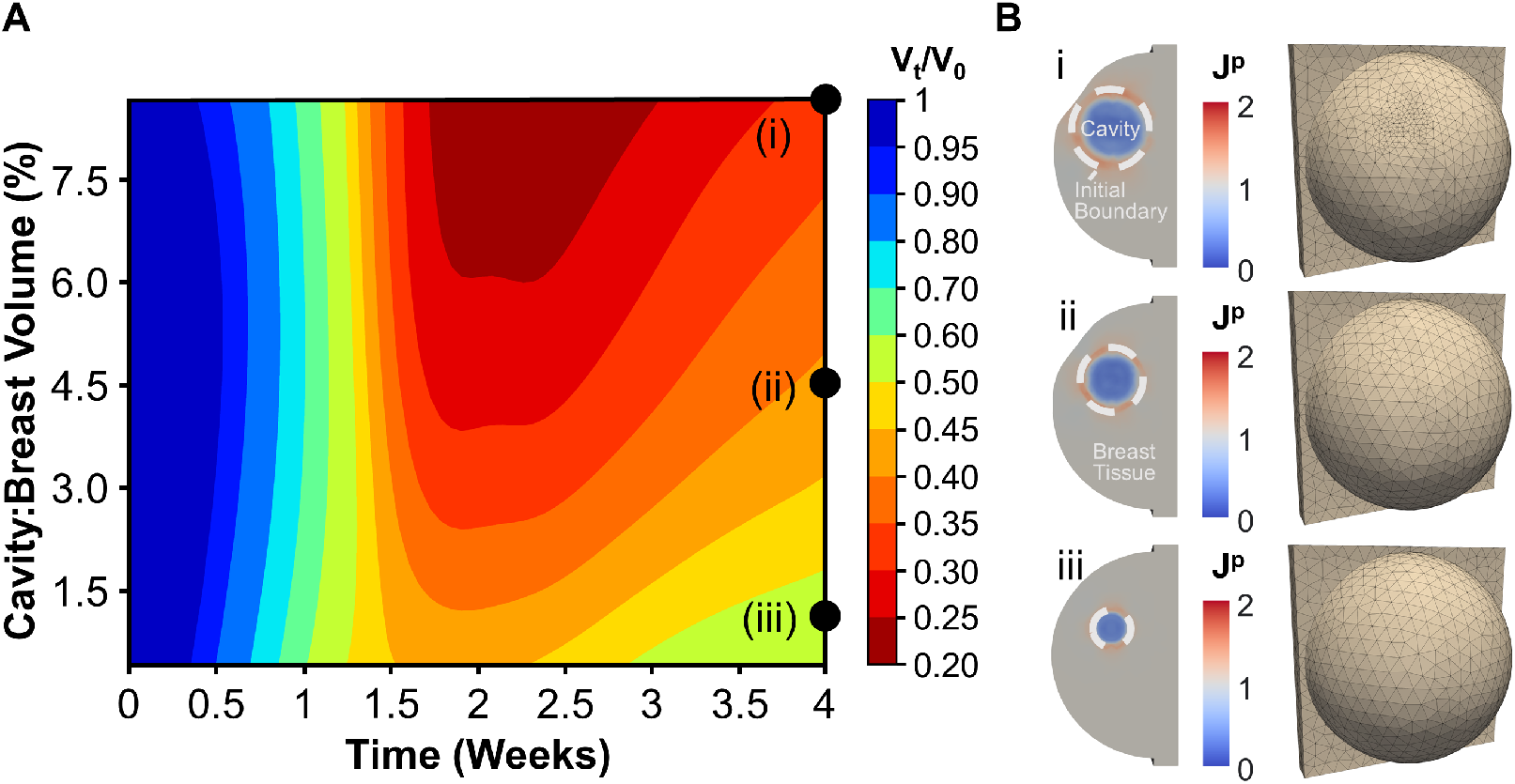
Effect of CBVP on cavity contraction and breast surface deformities. (A) Contour plot created with the re-calibrated GP model accounting for cavity volume as an input, which predicts time-dependent cavity contraction as a function of CBVP. (B) Simulations were run for three specific CBVP values to evaluate permanent tissue deformation *J*^*p*^ and breast surface deformation 4 weeks following lumpectomy for CBVPs of (i) 8.7%, (ii) 4.5%, (iii) and 1.0%.

Simulation results showed that smaller cavities contract at a faster rate compared to larger cavities, which is consistent with previously reported human wound contraction outcomes [55, 56]. Additionally, larger CBVP values showed a greater reduction in cavity volume (i.e., greater contracture). Finite element simulations were also conducted for specific CBVP values of i) 8.7%, ii) 4.5%, iii) 1.0% to verify accuracy of GP predictions and visualize breast deformations (Fig. 8B). As expected, permanent changes in breast volume and shape increased with cavity size, with similar permanent deformation values within the cavity centers (8B). Overall, larger breast surface deformation occurred with increasing CBVP. For instance, for a relatively small CBVP of 1.0%, there was no visible breast surface deformation 4 weeks post-surgery (8B). Increasing the CBVP to 4.5% resulted in moderate surface deformation, which became more severe for CBVP of 8.7% (8B). These results are consistent with reported clinical outcomes [57, 58, 21].

### 3.6. Effect of breast composition

To determine the effect of breast composition on BCS outcomes, the GP surrogate was further informed by running additional simulations including breast composition as an input variable. Specifically, recall that the material parameters *k*_0_, *k*_1_ were assigned based on the assumption of 70% adipose tissue and 30% fibroglandular tissue [38]. When evaluating the effect of breast composition, *k*_0_, *k*_1_ were modified according to the rule of mixtures by varying the percent of adipose to fibroglandular tissue. Following re-calibration, the GP model was used to predict 4-week post-surgical cavity contraction as a function of breast composition (Fig. 9.A). Clinically, breast composition is measured with the BI-RADS ranking system which reports the percentage of breast fibroglandular tissue [59]. As shown in 9A, cavities created in low density breasts (i.e., breasts consisting primarily of soft fatty tissue or scattered small regions of fibroglandular tissue) contracted more rapidly and to a greater extent than those in high density breasts (i.e., breasts consisting of heterogeneously or extremely dense fibroglandular tissue). Lower density breasts also gave rise to higher magnitudes of permanent contracture within the cavity, causing the surrounding breast tissue to be drawn in higher tension (Fig. 9B). Interestingly, permanent contracture was positively correlated with breast surface deformation, as lower breast densities were more prone to breast asymmetry (Fig. 9B). These results are consistent with clinical findings [18, 60, 19, 20].

**Figure 9:**
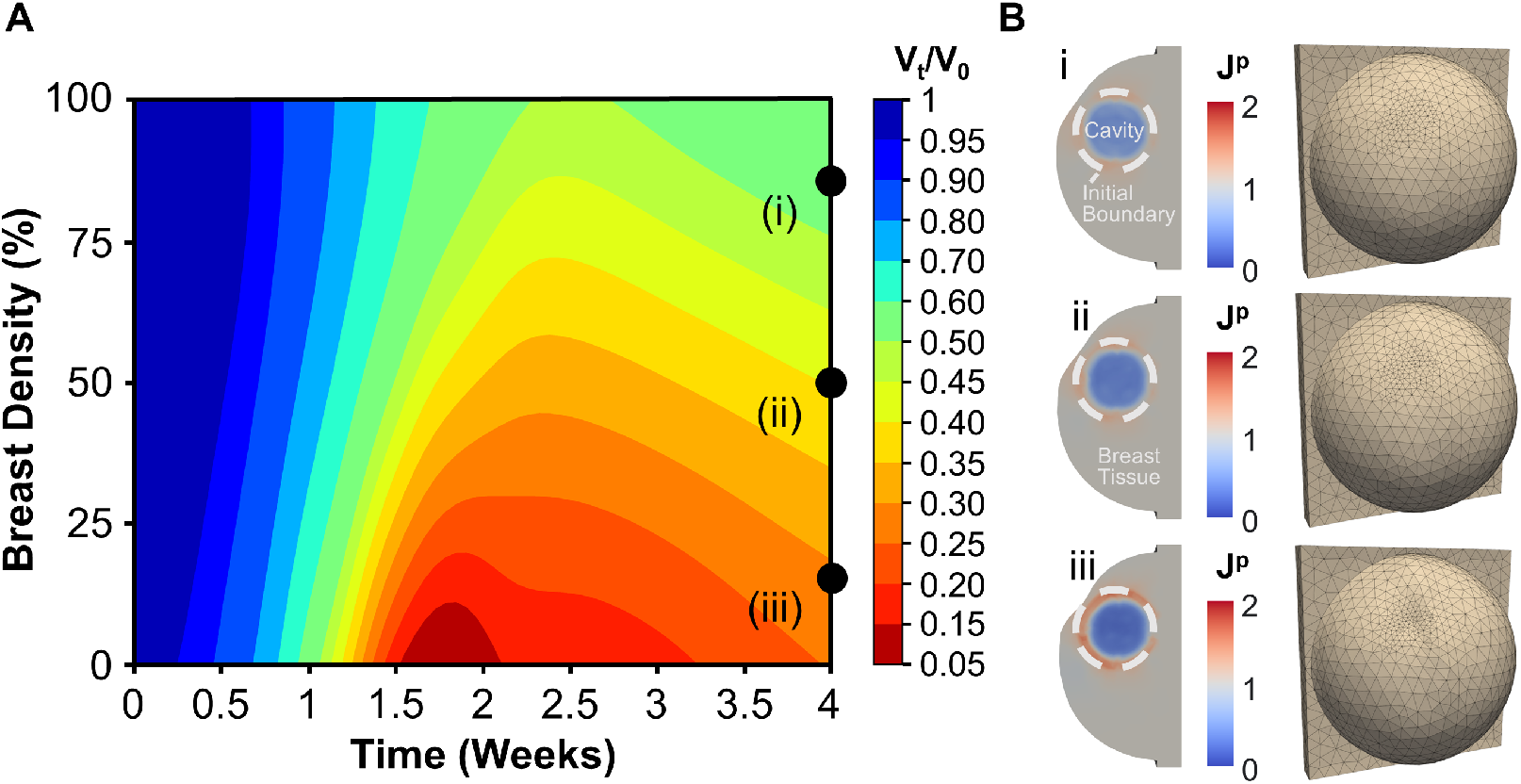
Effects of breast density on cavity contraction and breast surface deformities. (A) Contour plot created a recalibrated GP model taking into account breast composition as an input. (B) Finite element simulations for three breast densities show permanent tissue deformation *J*^*p*^ and breast surface deformation four weeks post-surgery for breast densities of (i) 85%, (ii) 50%, and (iii) 15%.

## 4. Discussion

Understanding the mechanobiology of breast cavity healing after lumpectomy is essential for improved prediction of post-surgical outcomes and individualized treatment planning for breast cancer patients. At present, there is a relatively high incidence of BCS-related breast deformities, with approximately one-third of women developing dents, distortions, and asymmetry between breasts [15, 18, 21, 49], which negatively impacts survivor self-esteem or quality of life [5]. While the significance of this problem has been recognized by the breast surgical community, there remains a fundamental lack of mechanistic and objective tools that define how various patient-to-patient factors affect post-surgical cavity healing and cosmetic outcomes. In this study, we developed a detailed finite element model of breast cavity healing after BCS that was calibrated using experimental porcine lumpectomy and previously published human clinical data. The computational model incorporated biological, microstructural, and mechanical variables that describe fundamental breast healing processes and relationships. The finite element model was designed to define how the coupling of mechanobiological cues and patient-specific breast characteristics (geometry, consistency, and biomechanics) contributes to temporal changes in cavity contraction and associated breast volume and surface deformations. Therefore, this model has the potential to help both surgeons and patients anticipate BCS healing and cosmetic outcomes.

Computational and mathematical descriptions of wound healing processes and outcomes have been a focus area of investigation for over three decades, with the majority of models describing cutaneous (skin) repair [24, 61]. The first wound healing model, proposed by Sherratt and Murray (1990) [62], did not consider mechanobiology or tissue mechanics when describing reepithelialization of skin. For this early model, activation and proliferation of epithelial cells was assumed to occur along a 1D wound in response to chemical cues. Such models have been refined over time to include more complex cellular and chemical reaction-transport phenomena associated with inflammation and angiogenesis [63, 64]. Increasing attention has also been given to fibroblast and myofibroblast activity and their impact on collagen deposition and remodeling [65, 66]. Coupling to nonlinear tissue mechanics has been explored extensively by our group and others in recent years [67, 68, 63, 64, 69, 23, 32, 30]. Specifically, our published models have leveraged prior modeling efforts and focused on adding detailed descriptions of local mechanobiological couplings between (myo)fibroblast activity and collagen remodeling to explain the observed macroscale changes in tissue mechanics and elastoplastic deformation. Our extensive work on the calibration of the 3D dermal model based on data from rat excisional wounds showed the model’s ability to predict a large set of experimental observations including treatment with collagen scaffolds, providing confidence in the fundamental relationships encoded in the model [30].

Here, we describe a finite element model of breast cavity healing following BCS that builds upon our previously published computational mechanobiological models of cutaneous wound healing [23, 32, 30]. At present, there are few models describing the healing of deep wounds, such as those associated with BCS, with the majority being adapted from early skin wound models. For example, with the goal of predicting wound healing following lumpectomy, Garbey and co-workers developed a 2D cellular automata model linked to a PDE describing cytokine signaling within skin wounds [27, 67]. Likewise, Vavourakis et al. adapted a finite element model of inflammation and angiogenesis initially introduced by Sherratt and Murray, coupling it with a finite element model of soft tissue biomechanics [29, 70]. In the present study, we modified our 3D dermal wound model [30] to include more realistic fibroblast migration, with dependence on both cytokine concentration and collagen density. We also implemented a generalized breast geometry that was based on human clinical data and adjusted tissue mechanical properties based on the literature. Biochemical and mechanobiological model parameters that were not well defined in the literature were tuned and optimized, allowing the computational model to be fit to experimental porcine lumpectomy data describing time-dependent changes in fibroblast migration and collagen deposition and human clinical data depicting the volumetric breast cavity changes that occur after BCS.

The calibrated model was designed to provide a new and useful tool for supporting future hypothesis generation, surgical visualization, and surgical decision-making. More specifically, we applied the model to define how patient-to-patient variability in breast and tumor characteristics affected breast contracture and breast surface deformation. When evaluating CBVP, model simulations predicted that larger cavities, specifically located within the outer quadrant of the breast, would contract more slowly but to a greater extent than smaller cavities. Additionally, as CBVP increased from 1.0% (13.24 *cm*^3^ volume; 2.94 cm diameter) to 8.7% (115.5 *cm*^3^; 6.04 cm diameter), resultant tissue permanent deformation profiles contributed to more severe breast distortions. These model predictions aligned well with previously published clinical perspectives that state that tumor size, breast tissue volume excised, and CBVP are major determinants of BCS cosmetic outcomes. Maximum tumor diameters between 2 cm and 4 cm are commonly used as selection criteria for BCS [71, 58]. Moreover, CBVP is highly correlated with breast cosmesis assessment scores and patient satisfaction following BCS. Specifically, more than 80% of women were very satisfied with breast aesthetic outcomes when their CBVP was less than 10% [57, 58, 21]. By contrast, CBVP greater than 20% led to high levels of patient dissatisfaction [57, 58, 21]. Tumor location is an important determinant of cosmetic outcomes and patient satisfaction following BCS, with proposed recommendations for maximum CBVP including the following: 18-19% for the upper-outer quadrant, 14-15% for the lower-outer quadrant, 8-9% for the upper-inner quadrant, and 9-10% for the lower-inner quadrant [16]. Such findings have led to proposed surgical decision-making algorithms, where breast volume, clinical tumor size, and tumor location serve as major determinants when choosing between breast surgical procedures to achieve satisfactory breast cosmesis and quality of life [15, 16]. While these algorithms are currently being evaluated in randomized controlled trials in patients who are candidates for both BCS and mastectomy, they do not account for mechanistic details of the wound healing response. As a result, they cannot predict breast deformation over time, account for further coupling phenomena such as individualized breast biomechanics, or aid in the design of new therapeutics.

The calibrated model was also used to determine how breast tissue density affected breast tissue contracture and breast shape following BCS. Human breasts, as well as other mammalian mammary glands, are composed of a heterogeneous mixture of fibroglandular and adipose tissue, which contributes to differences in consistency and biomechanical properties. Reported Young’s modulus ranges for human breasts vary from 0.7 to 66 kPa, depending on breast composition (e.g., percentage of fibroglandular to adipose tissue) [40, 47]. Model simulations evaluated breast densities representing 15% (*E*_*BT*_ = 14.5 kPa), 50% (*E*_*BT*_ = 25 kPa), and 85% (*E*_*BT*_ = 35.5 kPa), spanning the range of soft breast consisting primarily of fatty tissue to firm (stiff) breast consisting primarily of fibroglandular tissue. Our simulations predicted that cavities within low density, fatty breasts exhibit larger contracture compared to high-density, firm breasts. As a result, breast surface deformities were larger and more pronounced as breast density decreased. These results are in agreement with human clinical findings, as many studies have correlated through patient surveys and clinical analysis that patients with low breast density have higher chances of poor cosmetic results and low patient satisfaction after BCS [18, 60, 19, 20].

Mechanobiological parameters influencing cell contractility and plastic deformation were also proven to greatly impact cavity contracture and cosmetic outcomes. Through the sensitivity analysis shown in Figure 7, we were able to learn more about plausible parameter ranges and gain insight into complex parameter relationships. The parameters that were deemed to be the most sensitive to the mechanobiological response and contracture were *t*_*ρ*_ and *t*_*ρ,c*_. Therefore, it is important to ensure model accuracy regarding these two parameters. Both *t*_*ρ*_ and *t*_*ρ,c*_ were optimized based on clinical data evaluating time-dependent cavity volume changes. Compared to dermal wound healing models that considered fibroblast traction based on experimental evidence, our model’s optimized value for *t*_*ρ*_ was on the lower end of the established range [66, 67, 63, 64, 69, 32, 30]. Relative to the contractility of fibroblasts, the optimized *t*_*ρ,c*_ value for our model was also well within the broad range of values in other wound healing models [66, 67, 63, 64, 69, 32, 30]. To potentially reduce model uncertainty, future experimental studies could be conducted to measure and validate the contractile force of fibroblasts and myofibroblasts post-lumpectomy.

The present study was made possible by leveraging machine learning techniques to replace the high-fidelity computational model with inexpensive but accurate surrogates. In particular, GP surrogates were used to predict cell density, collagen density, and cavity contraction over time as a function of model parameters [72, 73, 74]. While a single simulation with the fully coupled model takes on the order of 20-72 hours to run (depending on model parameters), the GP evaluation can be performed in milliseconds. Therefore, this 10^7^ speed-up was crucial to perform the parameter optimization and sensitivity analysis. Although many machine learning techniques exist, the GP was applied due to its Bayesian construction which allows the estimation of both the desired quantity of interest and expected epistemic uncertainty (i.e., it provides an estimate of the confidence for a given prediction) [51]. This differentiates GP approaches from other popular tools such as artificial neural networks [51]. The prediction of the variance by the GP guided the selection of parameter combinations for which to evaluate the finite element model, akin to other active learning strategies using GPs [75].

The study is not without limitations. For the computational model, we implemented a generic human breast geometry that was informed through several clinical studies. Further, the model was calibrated by tuning mechanobiological parameters to fit clinical data of time-dependent cavity volume changes reported as an average of 34 patients. Future model iterations will incorporate more patient-specific data, which includes application of patient-specific breast geometries, tumor or cavity shapes and locations, and heterogeneous breast tissue compositions. Individual healing outcomes can then be compared to model predictions to further validate the model. Figure 10 shows an example of how the generalized human breast geometry can nonetheless be used to forecast possible poor cosmetic outcomes that patients may experience. The model also fails to incorporate other factors that can affect breast healing. For example, radiation therapy, which is commonly applied to patient breasts shortly after BCS, is not accounted for in the model. This is an area we hope to capture in future work. Addition of radiation therapy to the computational model would require changes cell death, inflammation, collagen deposition, and (myo)fibroblast contraction, ultimately leading to changes in mechanical properties and breast deformation. Although the mechanobiological model is able to accurately predict healing outcomes, the complexity of the model can be further expanded to include additional specific cellular players and processes such as neovascularization, various types of immune cells (e.g., macrophages or neutrophils), and edema related osmotic pressure and poroelastic response. Future model applications also include the design of therapeutic approaches (e.g., regenerative breast tissue fillers), enabling the promise of in silico trials for BCS before animals or human subjects are involved.

**Figure 10:**
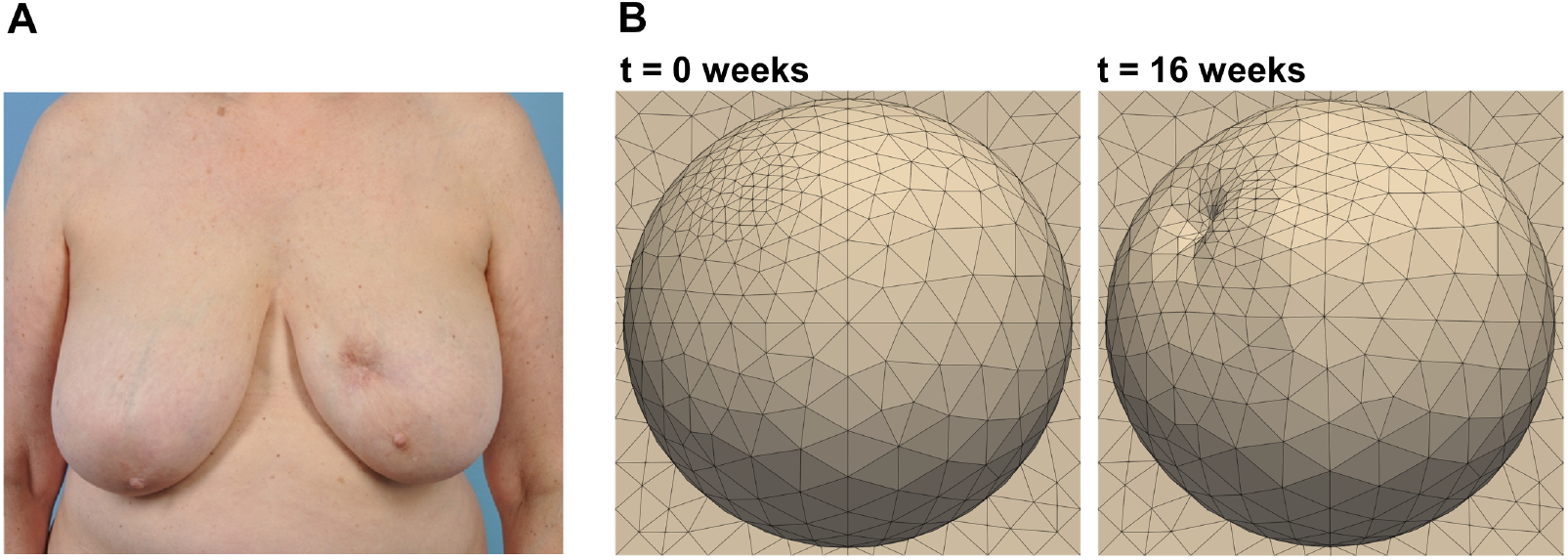
Comparison in the cosmetic outcomes after BCS between (A) a patient 5 years removed from BCS and (B) the generalized human breast geometry simulated 16 weeks post-surgery. (A) is reprised from Adamson et al. (2020) [76].

## 5. Conclusions

The presented computational model proved to effectively simulate the breast healing response following BCS, including fibroblast infiltration, collagen remodeling, and breast permanent deformation. Preclinical porcine data and human clinical data were used to inform time-dependent trends for fibroblast density, collagen density, and cavity volume change. The model was fit to this data by optimizing model parameters enabled by GP regression. Although previous models of wound healing after BCS have been developed, we advanced these efforts by implementing a detailed mechanobiological model coupled with the nonlinear mechanics of breast tissue, including large plastic deformation and collagen remodeling. Therefore, our model is uniquely suited for the prediction of scar tissue formation and breast deformation after BCS, which allowed us to gain insight into how key parameters and patientto-patient variability with respect to breast and tumor characteristics factor into the post-surgical cosmetic outcome. With this work presenting the foundation of the computational model, future efforts can be shifted to focus on patient-specific cases, addition of radiation therapy effects, and the design of therapeutic approaches (e.g., regenerative breast fillers).

## Acknowledgements

This work was supported, in part, by an NSF CMMI Multiscale Mechanobiology of Growth and Remodeling During Wound Healing grant (A.B.T.; 1911346). The preclinical porcine lumpectomy study was supported by an NSF SBIR Phase I award (S.V.-H.; 1913626). The authors acknowledge the Purdue University Histology Research Laboratory, a core facility of the NIH-funded Indiana Clinical and Translational Sciences Institute, for preparation of histological slides. Z.H., M.F., and C.M. were recipients of a Purdue Summer Undergraduate Research Fellowship (SURF). E.V. and D.S. are trainees of the NIGMS-funded Indiana Medical Scientist/Engineer Training Program (T32 GM077229) and the recipient of a NIDDK-funded predoctoral fellowship (T32 DK101000) and an Indiana CTSI predoctoral fellowship (UL1TR002529).

## Supplementary information

The finite element model is available in the following repository: https://github.com/zharbin/CBM_2023_BCS.

