## Supplementary Material for "Computational Mechanobiology Model Evaluating Healing of Postoperative Cavities Following Breast-Conserving Surgery"

| Parameter | Description | Value | Reference |
|---|---|---|---|
| $D_c$ $[mm^2/hr]$ | Cytokine Diffusion Coefficient | 0.01208 | [1, 2, 3] |
| $d_{\rho,c}$ $[mm^2/hr]$ | Cytokine-Increased Fibroblast Diffusivity | $6.12 \times 10^{-3}$ | Estimated |
| $d_{\rho,0}$ $[mm^2/hr]$ | Baseline Fibroblast Diffusivity | $6.12 \times 10^{-5}$ | [4] |
| $p_\rho$ $[1/hr]$ | Fibroblast Proliferation | $9 \times 10^{-4}$ | Estimated |
| $K_{\rho,c}$ [-] | Proliferation Saturation due to Cytokine | $1 \times 10^{-5}$ | [5] |
| $p_{\rho,e}$ $[1/hr]$ | Mechanoregulation of Fibroblast Proliferation | $p_\rho/2$ | [6] |
| $K_{\rho,\rho}$ [-] | Fibroblast Division Saturation | $550, 512.6$ | [7] |
| $d_\rho$ $[1/hr]$ | Fibroblast Death Rate | $p_\rho(1 - \rho_{phys}/K_{\rho\rho})$ | [7] |
| $p_{c,\rho}$ $[1/hr]$ | Fibroblast Secretion of Cytokine | $1.635 \times 10^{-18}$ | [5] |
| $p_{c,e}$ $[1/hr]$ | Mechanoregulation of Cytokine | $5.45 \times 10^{-18}$ | [5] |
| $K_{c,c}$ $[mol/mm^3]$ | Cytokine Saturation | 1 | [5] |
| $d_c$ $[1/hr]$ | Cytokine Death Rate | 0.005 | Estimated |
| $\rho_0$ $[cells/mm^3]$ | Nominal Fibroblast Density | 55051 | Estimated |
| $c_0$ $[g/mm^3]$ | Initial Cytokine Concentration Inside Cavity | $1 \times 10^{-4}$ | [5] |

Table 1: Parameters for the biochemical model. Parameters listed as estimated were selected in this work or modified from our previous wound healing models [5, 6].

| Parameter | Description | Value | Reference |
|---|---|---|---|
| $k_0$ $[MPa]$ | Linear Stiffness | $6.375 \times 10^{-3}$ | Estimated |
| $k_1$ $[MPa]$ | Compressibility | 0.317 | Estimated |
| $k_f$ $[MPa]$ | Fiber Stiffness | 0.015 | [8] |
| $k_2$ [-] | Nonlinear Stiffening | 0.048 | [8] |
| $\gamma_e$ [-] | Shape of Mechanosensing Curve | 5 | [5] |
| $\vartheta_e$ [-] | Midpoint of Mechanosensing Curve | 2 | [5, 9] |
| $K_{t,c}$ $[-]$ | Traction Saturation due to Cytokine | $1 \times 10^{-5}$ | [5] |
| $K_{\phi,c}$ $[-]$ | Collagen Production Saturation due to Cytokine | $1 \times 10^{-4}$ | [5] |
| $p_{\phi_e}$ $[1/hr]$ | Collagen Production Activated by Stretch | $p_\phi$ | [5] |
| $K_{\phi,\rho}$ $[-]$ | Collagen Production Saturation due to Collagen Fraction | $(\rho_0 * p_\phi)/d_\phi - 1$ | [5] |
| $d_\phi$ $[1/hr]$ | Collagen Degradation | $9.7 \times 10^{-4}$ | [10] |
| $d_{\phi,c}$ $[1/hr]$ | Collagen Degradation Activated by Cytokine | $8.81 \times 10^{-5}$ | [10] |
| $\tau_\omega$ $[hr]$ | Time Constant for Reorientation | $10/(K_{\phi,\rho} + 1)$ | [5] |
| $\tau_\kappa$ $[hr]$ | Time Constant for Dispersion | $1/(K_{\phi,\rho} + 1)$ | [5] |
| $\gamma_\kappa$ [-] | Shape of Dispersion Rate Curve | 2 | [5] |

Table 2: Parameters for the fully coupled mechanobiological model. Parameters listed as estimated were selected in this work or modified from our previous wound healing model [5, 6].

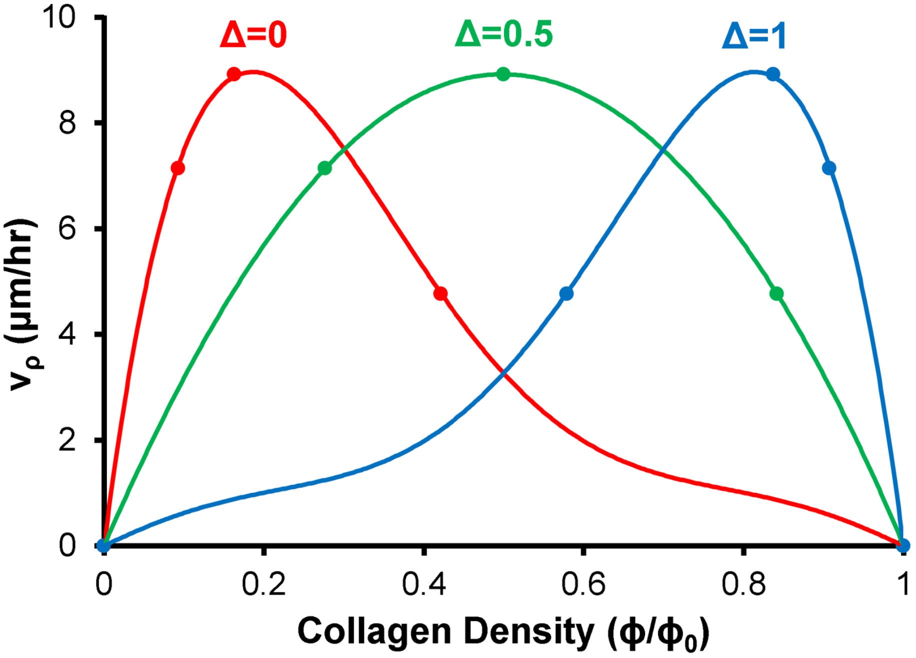


Figure 1: Fibroblast speed with respect to collagen density ($v_\rho(\phi)$) and its dependency on $\Delta$. Example $v_\rho(\phi)$ curves are shown with $\Delta$= 0 (red), 0.5 (green), 1 (blue). The function $v_\rho(\phi)$ was initially informed through [11, 12] through 5 data points displayed on the line $\Delta = 0$ while assuming $v_\rho(\phi) = 0$ for $\phi$= 0 and 1. Due to the limited data and uncertainty, parameter $\Delta$ was created to shift the 5 data points and skew the interpolated function. $\Delta$ was further investigated in the biochemical GP, where it was determined that the optimum value was $\Delta = 0$.

### Histological Image Analysis Methodology

#### Quantifying Fibroblast Density

1. Count Red Blood Cells (RBC)
   (a) Adjust Color Balance ([Minimum, Maximum])
      i. Red: [0,0]
      ii. Green: [0,100]
      iii. Blue: [0,0]
   (b) Convert Image Type From RGB Color to 32-bit
   (c) Apply Threshold ([0,$\sim$ 220])
   (d) Apply Watershed Segmentation
   (e) Analyze Particles for RBC Count
      i. Cell Size ([Minimum, Maximum]): [4,$\infty$]
2. Count All Cells
   (a) Adjust Color Balance ([Minimum, Maximum])
      i. Red: [70,220]
      ii. Green: [0,0]
      iii. Blue: [0,0]
   (b) Convert Image Type From RGB Color to 32-bit
   (c) Apply Threshold ([0,$\sim$ 245])
   (d) Apply Watershed Segmentation
   (e) Analyze Particles for All Cell Count
      i. Under "Set Measurements" Select Original Slide Under "Redirect to:"
      ii. Cell Size ([Minimum, Maximum]): [4,$\infty$]
   (f) Evaluate Modal Gray Value ($\leq\sim$125) to Isolate and Count Immune Cells
3. Fibroblast Count = All Cell Count - RBC Count - Immune Cell Count

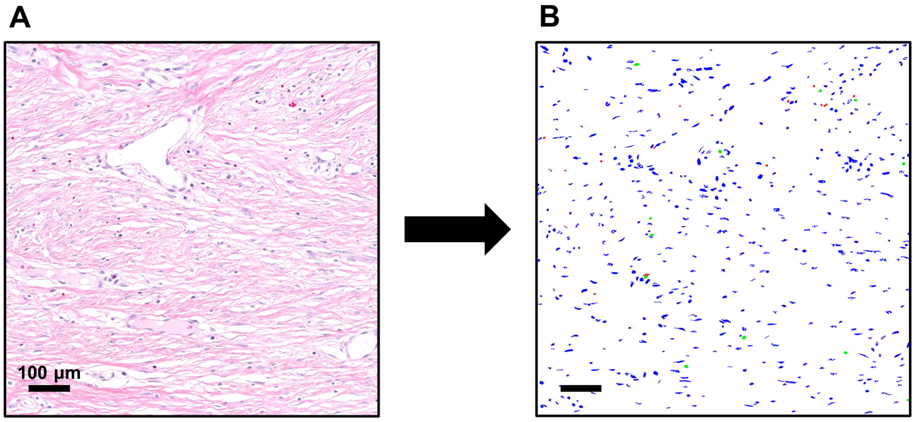


Figure 2: Result of post-processing individual regions (500 x 500 $\mu$m$^2$) obtained from porcine lumpectomy histology slides. (A) Regions were captured through Aperio ImageScope sampling across the entire cavity domain. Pictured is an example region from a histology slide 16 weeks post-surgery. (B) Using the procedures described above, regions were processed in ImageJ to quantify the number of fibroblasts (blue), red blood cells (red), and immune cells (green).

#### Calculating Collagen Density

1. In the 500 x 500 $\mu$m$^2$ histology region, select a small rectangular area ($\sim$100 $\mu$m$^2$) that contains no cells.

2. Measure for average pixel intensity in the small rectangular area. Note: Pixel intensity varies between 0 (black) and 255 (white).

3. Repeat steps 1 and 2 for a 500 x 500 $\mu$m$^2$ histology region that contains healthy breast connective tissue.

4. Calculate the estimated collagen density through the following equation:

$$(\phi/\phi_0)_{est.} = \frac{I_{scar}-255}{0.3*(I_{connective}-255)}$$

where $I_{scar}$ is the intensity of the scar tissue at the analyzed week and $I_{connective}$ is the intensity of the connective tissue.
